# Leafhoppers’ pest status in Québec from 1868 to 2025

**DOI:** 10.64898/2026.05.26.727661

**Authors:** Abraão Almeida Santos, Edel Pérez-López

## Abstract

Over the past decade, leafhoppers (Hemiptera: Cicadellidae) have gained increasing attention as pests and vectors of phytoplasma diseases in Québec.
Drawing on historical and modern reports spanning 1868 to 2025, we investigate whether leafhopper pest status has been changing in the province, focusing on two leafhopper groups: those that overwinter in Québec (local species) and those that do not (migratory species).
Local species have been reported as pests since at least 1886 and have been considered of secondary significance. Migratory species, first mentioned in 1908, have been listed more frequently as economically important.
*Empoasca fabae* and *Macrosteles quadrilineatus* were associated with 10 crops, accounting for 11–88% of adult specimens in reported crop assemblages. On the other hand, local species, such as *Erythroneura* ssp. and *Zonocyba* (= *Typhlocyba*) *pomaria*, were reported mainly on grapevines and apples, respectively.
We found that the first field reports of leafhoppers from 1997 to 2025 are advancing in crops where migratory species are most frequently reported (e.g., potato and strawberry), whereas in crops where local species appear to predominate, no significant association was found (e.g., apple and grapevine).
Our findings indicate that recent changes in leafhopper pest status in Québec are likely due to migratory species, a group that has historically been a significant component of the leafhopper assemblage on agricultural crops. In Québec agriculture, what was once ‘*loin des yeux, loin du cœur*’ (out of sight, out of mind) appears to be shifting toward ‘*près des yeux, près du cœur*’ (now visible and therefore important).

**GRAPHICAL ABSTRACT AND HIGHLIGHTS:** 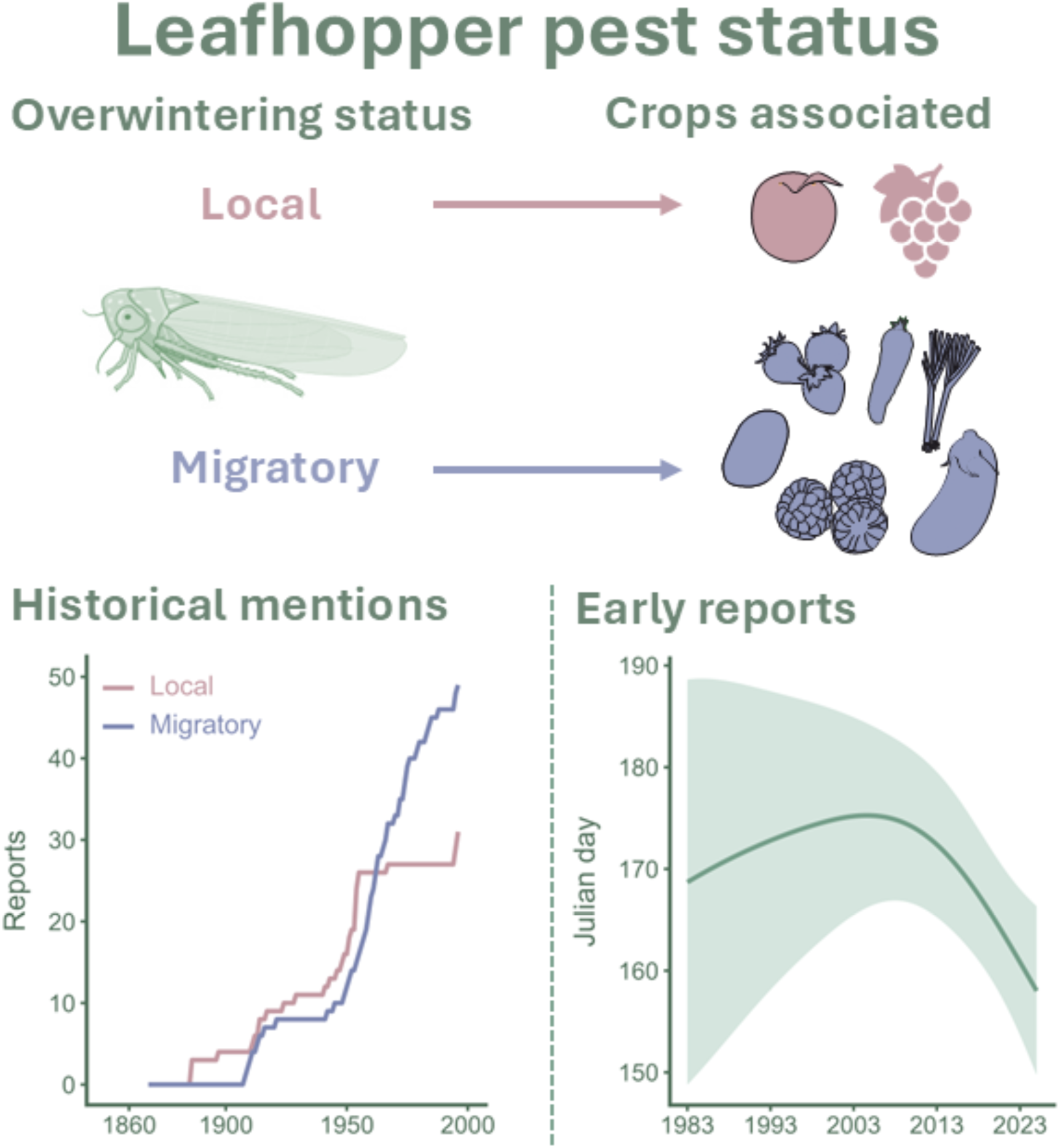

- The pest status of leafhoppers has changed in Québec over recent decades, but whether local or migratory species are driving this shift remains unknown.
- Local species have been recognized as pests since at least 1886, and migratory species since 1908. However, migratory species were more frequently reported in historical documents.
- Migratory species (*Empoasca fabae* and *Macrosteles quadrilineatus*) appear to be driving the recent status change, as evidenced by earlier first reports and presence across multiple crops.

## INTRODUCTION

The French expression ‘*Loin des yeux, loin du cœur*’ (out of sight, out of mind) captures a fundamental concept about how we perceive insect pest species: when they are neither easily noticeable nor associated with economic losses, they are often disregarded, and their perceived relevance fades. Leafhoppers (Hemiptera: Cicadellidae) exemplify this dynamic. These insects are recognized vectors of multiple plant pathogens (Olivier et al., 2009; Weintraub & Beanland, 2006) and among the reasons why they are getting into growers and decision makers’ radars are (i) due to the detection of an invasive vector species (Faris et al., 2024; Parent et al., 2019); (ii) outbreak populations causing economic losses (Machado et al., 2024; Moritz et al., 2025); and (iii) increased leafhopper-transmitted disease incidence (Brochu et al., 2023; Duffeck et al., 2025). However, outside these triggering conditions, when they are ‘*loin des yeux*’, they remain ‘*loin du cœur*’.

Perceived significance of leafhoppers as pests varies considerably: outbreaks, new detections, and epidemic conditions attract considerable attention, whereas periods of relative inactivity make them less of an economic concern and consequently decrease research focus (Ross, 1952). Nonetheless, such episodic attention poses significant challenges for understanding, predicting, and managing leafhopper pests, especially in northern agricultural areas amid ongoing warming. The northward shift of pests and pathogens has been documented in response to warming climates (Bebber et al., 2013, 2014), leading, in some cases, to earlier detection and increased crop damage (Baker et al., 2015; Plante et al., 2024). However, the relationship between these shifts and warming conditions depends on the specific temporal and research contexts in which leafhopper species are studied. This complexity complicates efforts to distinguish true population trends from potential biases in surveillance or reporting (Didham et al., 2020; Lehmann et al., 2020).

Québec Province, in Eastern Canada, offers a compelling case for examining these dynamics among leafhopper pest species. In this province, the earliest formal records of leafhoppers affecting crops date back to the late 19th century, with the first descriptions of Hemiptera species by Léon Provancher (1886). Subsequently, species were referenced in annual survey reports when their presence on crops was considered as somehow economically important (Arcelin & Kushalappa, 1991; Hudon & Martel, 1973; Macnay, 1958, 1961; Maheux & Petch, 1929; Petch, 1913). During the period from 1950 to 1970, characterized by a high incidence of phytoplasma disease reports in vegetable and small fruit crops (Chiykowski et al., 1973; Miller & De Lyzer, 1960), intermittent annual monitoring was conducted in 1959, 1960, 1961, and 1964 (Lachance et al., 1961; Perron & Crête, 1968). Aside from this period, leafhoppers fade from the economic perspective on pest groups affecting crops in Québec and, consequently, from the survey focus.

However, in the past decade, they have again gained increasing attention as pests in multiple crops, including as vectors of phytoplasma diseases. Outbreaks in vegetable crops in 2016 prompted continuous monitoring (RAP, 2025), the establishment of insecticide efficacy and application thresholds in crops where pressure was considered high (Khiari, 2020; Martinez et al., 2022), including the integration of multiple control methods (Shi et al., 2025). Concurrently, reports from 2012 to 2022 documented phytoplasma diseases affecting 34 plant hosts in the province, including new phytoplasma-host strain records and an apparent re-emergence of diseases that had been widespread between 1950 and 1970 (Brochu et al., 2023). Additionally, the detection of economically significant diseases in Canada, such as *Bois noir* (Rott et al., 2007), prompted surveys of phytoplasma prevalence in grapevines and leafhopper species, confirmed or suspected to be vectors, across the country, including in Québec (Olivier et al., 2014; Saguez et al., 2014).

Nonetheless, such patterns are not without historical precedent in Eastern Canada. Miller (1956) reported an apparent shift in leafhopper pest status in vegetable crops in Southwestern Ontario, linked to migratory species (i.e., species unable to overwinter in Canada), and attributed recurring outbreaks to a combination of a more favorable climate, high crop diversity, and agricultural intensification. Miller already observed that such rapid changes were especially difficult to interpret or to attribute to specific causes, given the absence of continuous population monitoring (Miller, 1956). Importantly, this distinction between migratory and local leafhopper pest species is particularly relevant for Québec. Migratory species, whose annual arrival depends on wind patterns and on source populations further south, are likely more responsive to climate variability and may show earlier spring detections under warming conditions (Baker et al., 2015; Carlson et al., 1992; Jacques et al., 2025; Plante et al., 2024; Santos et al., 2025a; Sidumo et al., 2005; Wallis, 1962). In contrast, local species that overwinter in the province may exhibit more stable phenology and be less sensitive to interannual climate fluctuations (Bostanian et al., 2006; Plante et al., 2024). Some evidence supports this view, as rising temperatures have advanced the phenology of migratory leafhopper species in North America, contributing to earlier arrivals and increased crop damage (Baker et al., 2015; Maredia et al., 1998; Plante et al., 2024).

If the pest status of leafhoppers has changed in Québec, an early sign would be earlier detection of migratory species, although local species may also contribute through shifts in seasonal activity or abundance. However, because historical records are shaped by episodic attention, a single data source is insufficient to detect sustained changes in pest status. We therefore used multiple approaches to evaluate our hypothesis, integrating data from a modern phytosanitary surveillance network with historical records spanning 1868 to 2025. Specifically, we addressed three questions: (i) Are first field reports of leafhoppers appearing earlier? (ii) Do these early shifts indicate changes in pest status? (iii) Are these changes associated with migratory species, local species, or both?

## METHODS

In this study, we integrated multiple data sources to reconstruct changes in leafhopper pest status in Québec: reports providing systematic seasonal and first-record data; historical documents and museum specimens detailing long-term species presence and pest-crop associations; and published assemblage surveys documenting species composition and relative abundance across crops. This approach allowed us to identify shifts in pest status and evaluate the roles of migratory and local species on these shifts.

### Data collection and criteria for first reports

To assess whether the first reports of leafhoppers in agricultural systems in Québec are advancing, we conducted a systematic search of historical and scientific documents with no date restrictions, following the principles of systematic review adapted to archival research. All searches focused on documents in English or French, considering that Québec is a francophone province in Canada. We applied two inclusion criteria: (i) the work must have been conducted within Québec or include Québec as a study location, and (ii) the document must include seasonal leafhopper sampling or reporting under field conditions, with the first detection date sufficiently precise to assign a Julian day. Documents presenting only aggregated data (e.g., monthly or annual summaries) were excluded.

### Scientific literature

In January 2026, we searched the Web of Science Core Collection using the string: ‘leafhopper’ OR ‘cicadellid’ OR ‘Cicadellidae’ AND ‘Canada’. This yielded 201 unique documents. Titles and abstracts were screened against the inclusion criteria, followed by full-text assessment of potentially eligible studies.

### Phytosanitary reports

The primary source for current pest and disease field reports in Québec is the ‘*Réseau d’avertissements phytosanitaires*’ (RAP; https://www.agrireseau.net/rap?a=1Csort=2), which has published weekly bulletins since 2004, mainly during the growing season (May to October), covering over 20 agricultural crops. We searched all RAP bulletins from 2004 to 2025 using the French keyword string ‘cicadelle’ OR ‘cicadelles’. For each year, we recorded the date of the first report mentioning leafhopper presence for each crop. When no specific date was provided, we used the bulletin publication date as a proxy, introducing a potential offset of up to 7 days. From RAP reports, we also compiled a list of leafhopper species and their associations with phytoplasma diseases from 2004 to 2025.

### Historical documentation

Because historical documents are not indexed in electronic databases and many exist only in print, we conducted targeted archival searches using the Biodiversity Heritage Library (BHL; https://www.biodiversitylibrary.org/) and other sources to trace associations between leafhopper pest species and crop field reports in Québec. These documents provide context on the economic importance of leafhoppers over time.

For each document, we reviewed the title, abstract, and species index when available, then searched for French and English keywords: aster yellow, *cicadelle*(s), leaf-hopper(s), leafhopper(s), hopper(s), green petal, *jaunisse de l’aster*, phytoplasmas, virus-like, yellows, tip-burn, and hopperburn. When mentions were found, we extracted the year, leafhopper species, associated crops, and diseases. All leafhopper species names were verified for synonyms using the World Auchenorrhyncha database (Dmitriev et al., 2026).

We examined the following historical sources (URLs and access details are provided in **Supporting Material S1**): *Le Naturaliste Canadien* (1868–2025); Provancher Hemiptera checklist (1886); the Journal of the Entomological Society of Ontario (1871–2025); publications of the *Société d’entomologie du Québec* (1956–2021); the *Rapport annuel de la Société de Québec pour la protection des plantes* (1908–1961); Phytoprotection (1963–2025); the *Rapport annuel de la Société de Pomologie et de Culture Fruitière* (1894–1960); A Synopsis of Economic Entomology (1914); *Les principales espèces d’insectes nuisibles et les maladies végétales* (1916); the bulletin *Nouv’Ailes* (1999–2023); and the Canadian Plant Disease Survey Archive (1920–2025).

### Museum records

For each leafhopper species identified in the RAP network and historical documents, we searched for specimens collected in Québec, recording the collection year and Julian day. We obtained records from the Canadian National Collection of Insects, Arachnids, and Nematodes (CNC, Ottawa) via its online portal (https://cnc.agr.gc.ca/taxonomy/TaxonMain.php). We also searched the Hemiptera collections of the Insectarium de Montréal, the Ouellet-Robert Entomological Collection (Université de Montréal), and the Lyman Entomological Museum (McGill University) using the Canadensys portal (https://www.canadensys.net/) but found no leafhopper records for the target species in any of these collections. The René-Martineau Insectarium collection yielded seven records of *Empoasca fabae* (Harris) (1933–1934) without collection dates. We also verified the Provancher Hemiptera collection at Université Laval in March 2026, but specimens lacked collection dates.

### Leafhopper assemblage on crops

To contextualize the relative abundance of species within the leafhopper assemblage, we examined published leafhopper surveys conducted in Québec crops identified through our literature and historical search. Studies were included if they provided quantitative data (e.g., specimen counts). From each study, we extracted adult specimen counts and classified each species as either migratory (annual immigrants unable to overwinter in Québec) or local (overwintering in Québec) based on published biological data. Two species, *Empoasca fabae* (Baker et al., 2015; Carlson et al., 1992) and *Macrosteles quadrilineatus* (Forbes) (Wallis, 1962) are well-documented as migratory in the province. All other species were classified as local, but we acknowledge that limited bioecological data on these species introduce uncertainty in this classification. For each study, we calculated the proportion of specimens belonging to migratory versus local species.

### Statistical analyses

To test whether leafhopper reports are occurring earlier over time while allowing for nonlinear trends, we modeled the Julian day of the first report (DOY) as a function of year using a generalized additive mixed model of the form: DOY ∼ s(Year, k = 5) + s(Crop, bs = “re”), with a first-order autoregressive [AR(1)] correlation structure: corAR1(form = ∼ Year | Crop). We included the year as a smooth term and the crop as a random intercept. The estimated autocorrelation parameter (Φ = 0.60, 95% CI: 0.39–0.74) indicated moderate temporal dependence. Model diagnostics (normalized residual autocorrelation, homogeneity of variance, normality) showed no significant autocorrelation remained (Box-Ljung test: lag 5, χ² = 6.04, p = 0.302; lag 10, χ² = 14.19, p = 0.165). Residual diagnostic plots are available in **Supporting Figure S1**.

To complement the overall trend and assess crop-specific rates of change, we fitted individual linear regression models for crops with at least five years of data, with year as the predictor and the first report date (Julian day) as the response. Model assumptions were verified, and for the potato data, we report heteroscedasticity-consistent standard errors. The estimated slope coefficients for each crop, expressed as the annual rate of change in days per year, along with their 95% CIs, were displayed in a forest plot. All analyses were conducted in R (version 4.4.2, R Core Team, 2025) using packages mgcv (v. 1.9-4) (Wood, 2026), lme4 (v. 1.1-36) (Bates et al., 2015), and performance (v.0.15.3) (Lüdecke et al., 2021). Plots were generated using ggplot2 (v.4.0.1) (Wickham, 2016).

## RESULTS

### First reports survey summary

Our survey yielded 151 records of first leafhopper reports across 16 crops from 1960 to 2025 (**Supporting Table S1**). The RAP network contributed the most records (141), with an additional 10 records from literature (Bostanian et al., 2003; Gagné et al., 1985; Jacques et al., 2025; Lachance et al., 1961; Plante et al., 2024; Shi et al., 2025).

The earliest seasonal record, dated 9 June 1960, for alfalfa (Lachance et al., 1961), was excluded from our analysis due to the 23-year gap until the next observation in 1983. The first report included in our analysis was from 1983 (Gagné et al., 1985), with only three additional records appearing before 2000 (**Figure 1a**). Records increased substantially after the establishment of the RAP network in 2004. Apple, lettuce, grapevines, and potato accounted for the most records (≥ 19 each), while blueberry, cabbage, maple, onion (categorized in the RAP network with garlic and leek), and timothy had the fewest (≤ 3 each) (**Figure 1a**).

**FIGURE 1.**
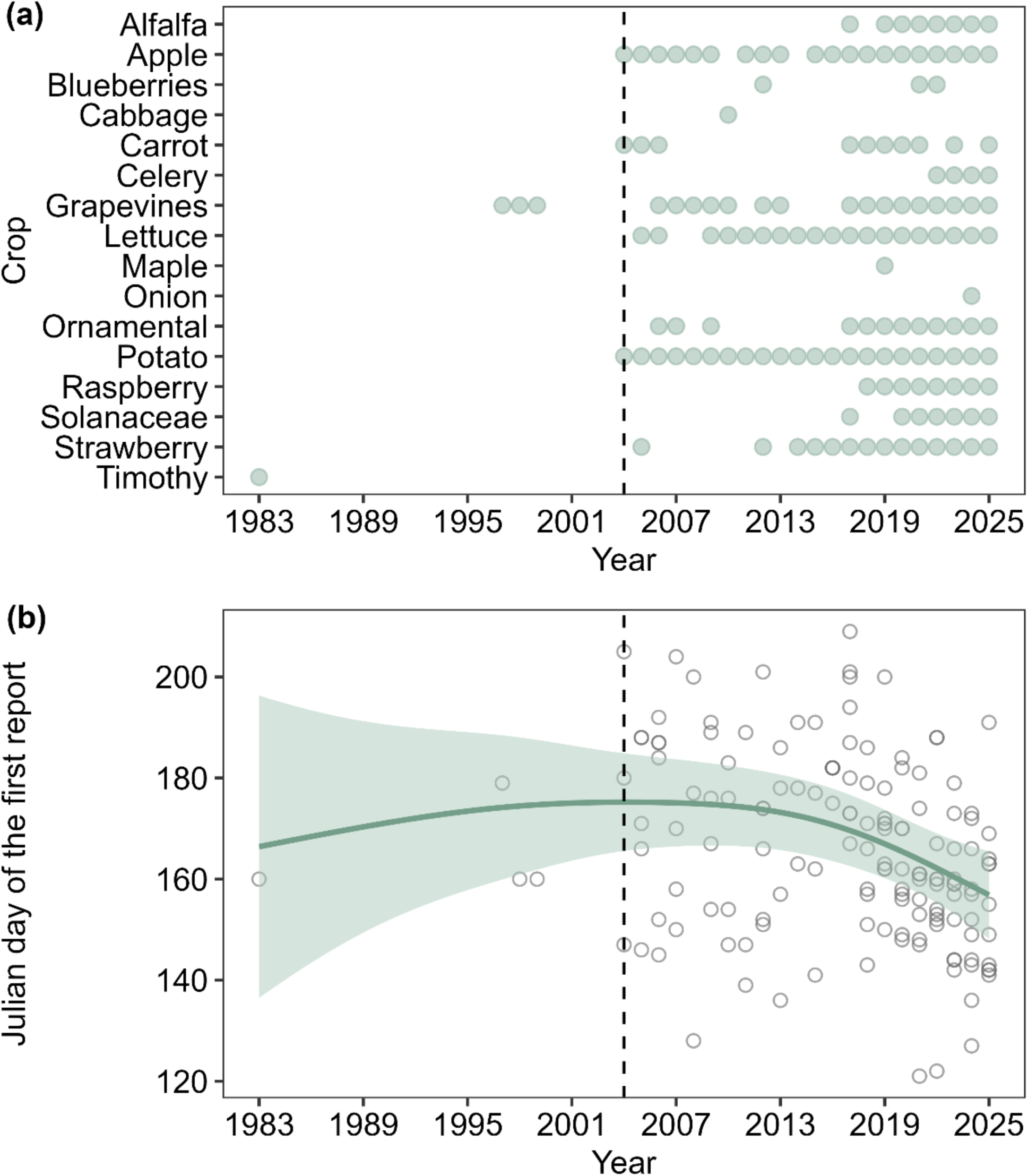
**(a)** Distribution of leafhopper first reports across 16 crops in Québec (Canada), from 1983 to 2025 (n = 150). **(b)** The relationship between the first report date (Julian day) and the year. The dark green line shows the trend fitted by a generalized additive mixed model, with shading representing the 95% confidence interval (see Table 1 for model details). The vertical gray dashed line in **(a)** and **(b)** marks the start of the RAP network in 2004. Data are available in **Supporting Table S1**.

**TABLE 1.**
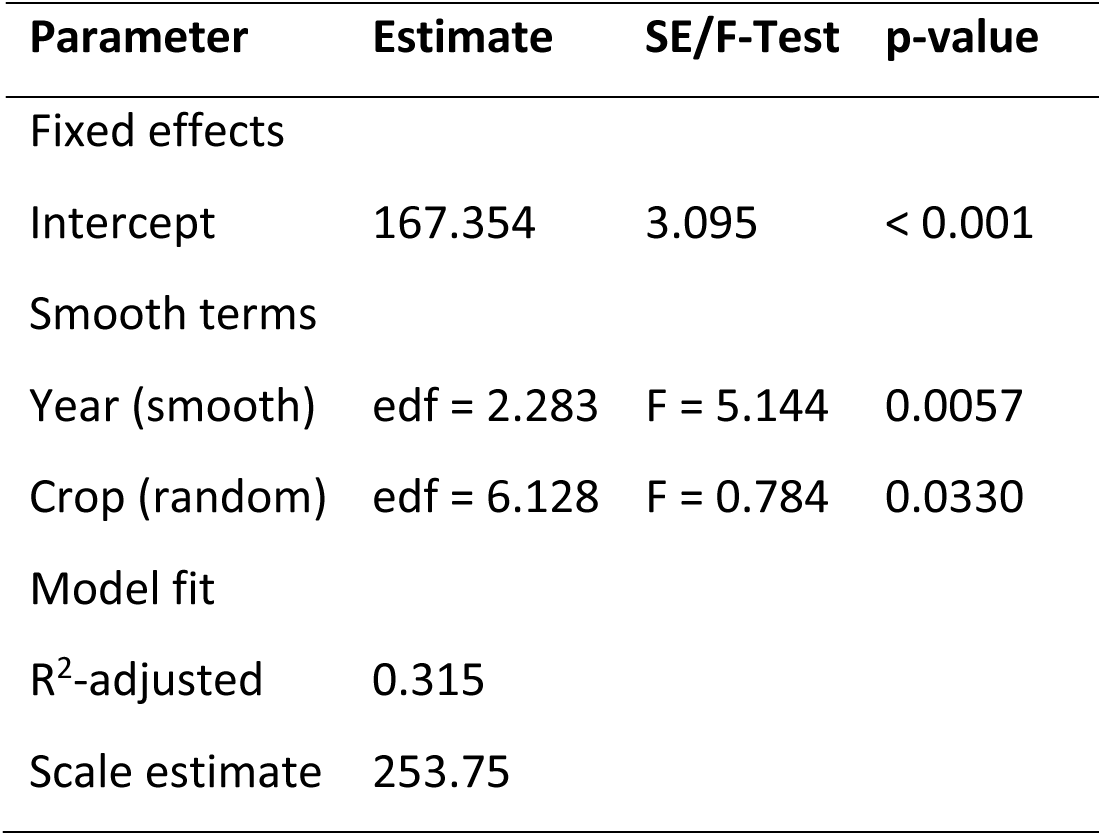
Summary of the generalized additive mixed model for the first report of leafhoppers (Julian day) over the years (1983-2025) across 16 crops (n = 150). edf = effective degrees of freedom.

### Leafhopper field reports are occurring earlier over time

First leafhopper reports showed a significant nonlinear trend over the 1983–2025 period (**Figure 1b**, **Table 1**). During the pre-2000 period, data were too sparse to detect a directional trend. First report dates were the latest around 2013, after which they declined steadily through 2025 (**Figure 1b**). The smooth term for year was significant (*p* = 0.0057), and the model explained 31.5% of the variance, with crops accounting for substantial random variation (**Table 1**).

### Crop-specific trends in first report dates

Significant negative associations (*p* < 0.05) between first report date and year were detected for five of the ten crops: carrot, lettuce, ornamental nurseries, potato, and strawberry (**Figure 2**). First reports advanced most rapidly in strawberry (–3.48 days per year), while in carrot, lettuce, potato, and ornamental nurseries, the advance was approximately 2 days per year (**Figure 3**). No significant associations were found for the remaining five crops: alfalfa, apple, grapevines, raspberry, and solanaceous crops (eggplant, tomato, pepper) (**Figures 2** and **Figure 3**).

**FIGURE 2.**
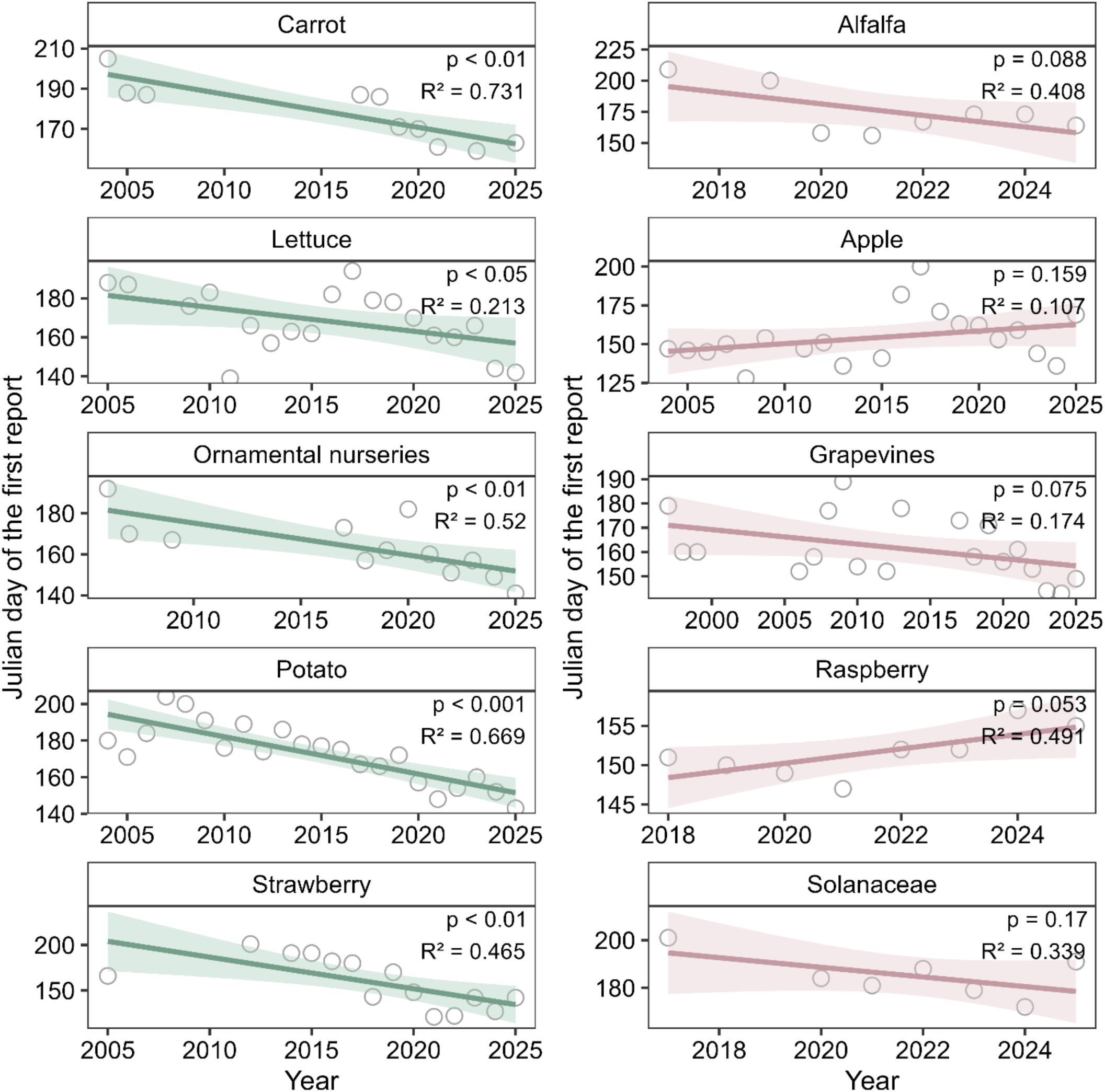
Relationship between first leafhopper report date (Julian day) and year for ten crops in Québec (Canada) with sufficient data (n ≥ 5 reports). Crops are arranged with significant associations (p < 0.05) on the left and non-significant associations (p ≥ 0.05) on the right. A separate linear model was fitted for each crop, and significant associations are shown in dark green; non-significant associations are shown in dark pink. Shading indicates 95% confidence intervals. Model significance (p-value) and coefficient of determination (R²) are displayed within each crop panel.

**FIGURE 3.**
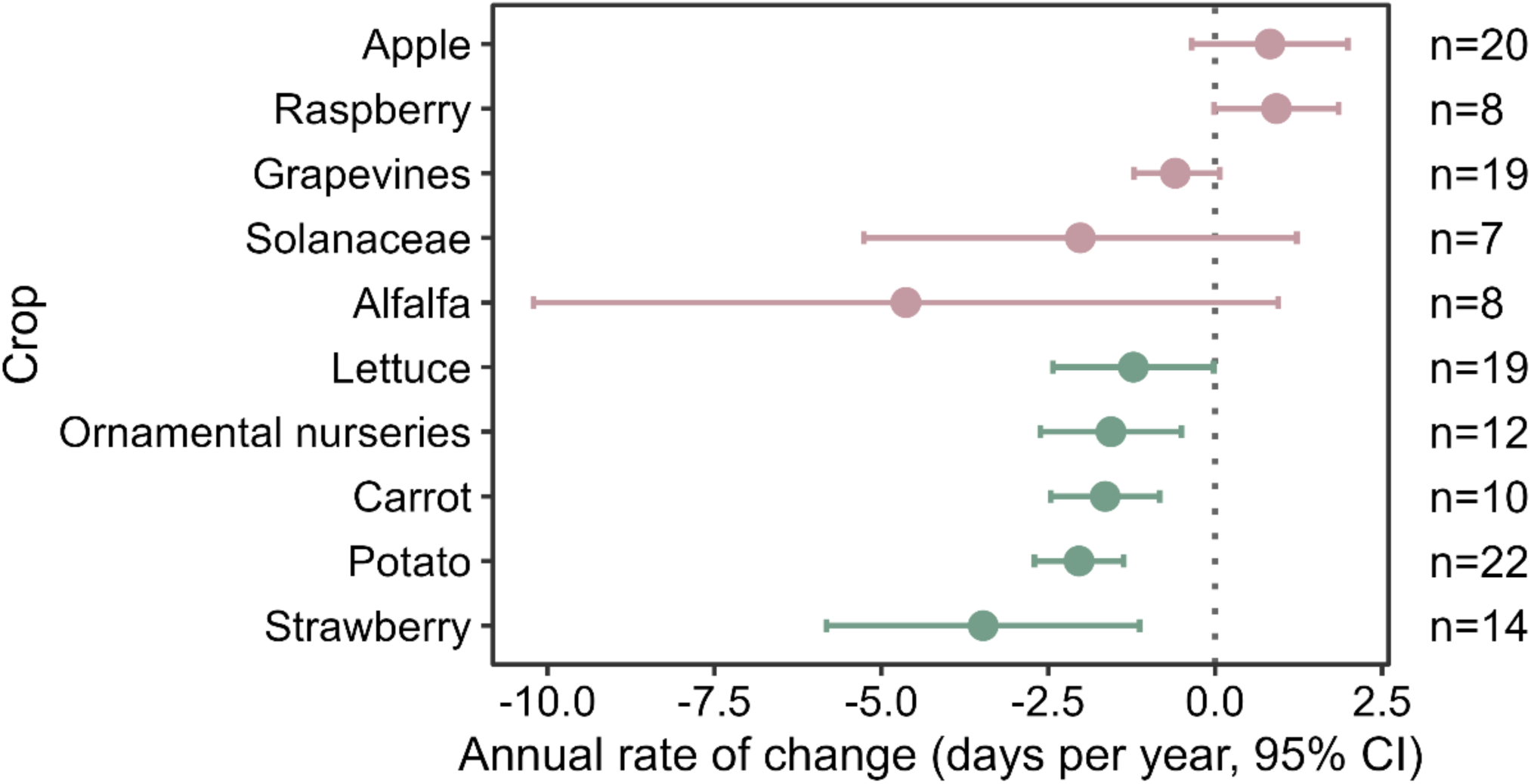
Estimated annual rate of change (days per year) in the first report date of leafhoppers for ten crops in Québec (Canada) with sufficient data (n ≥ 5 reports). Points represent the year effect (slope) from linear regression models fitted to each crop (Figure 3), with 95% confidence intervals (horizontal lines). Negative values indicate earlier first reports. Crops with confidence intervals that do not cross zero (dashed vertical line) show statistically significant trends (p < 0.05). Crops are ordered from those with earlier first reports (dark green) to those with later first reports (dark pink). Sample sizes (n) are shown next to the corresponding crop labels.

### Leafhopper species and phytoplasma disease reports in the RAP network

Seven leafhopper species were mentioned in RAP network reports between 2004 and 2025: *Empoasca fabae*, *Erythroneura comes* (Say), *Erythroneura tricincta* (Fitch), *Erythroneura vitis* (Harris), *Macrosteles quadrilineatus*, *Ophiola corniculus* (Marshall), and *Zonocyba* (= *Typhlocyba*) *pomaria* (McAtee) (**Table 2**). The two migratory species were reported across multiple crops, with *E. fabae* documented in 10 crops. The remaining five species are considered local in Québec and were reported on unique crops: the three *Erythroneura* species on grapevines, *O. corniculus* on highbush blueberry, and *Z. pomaria* on apple. In ornamental nurseries, initial reports mentioned leafhoppers only to the genus (*Empoasca*, *Graphocephala*, *Macrosteles*), with later reports specifying *E. fabae*.

**TABLE 2.**
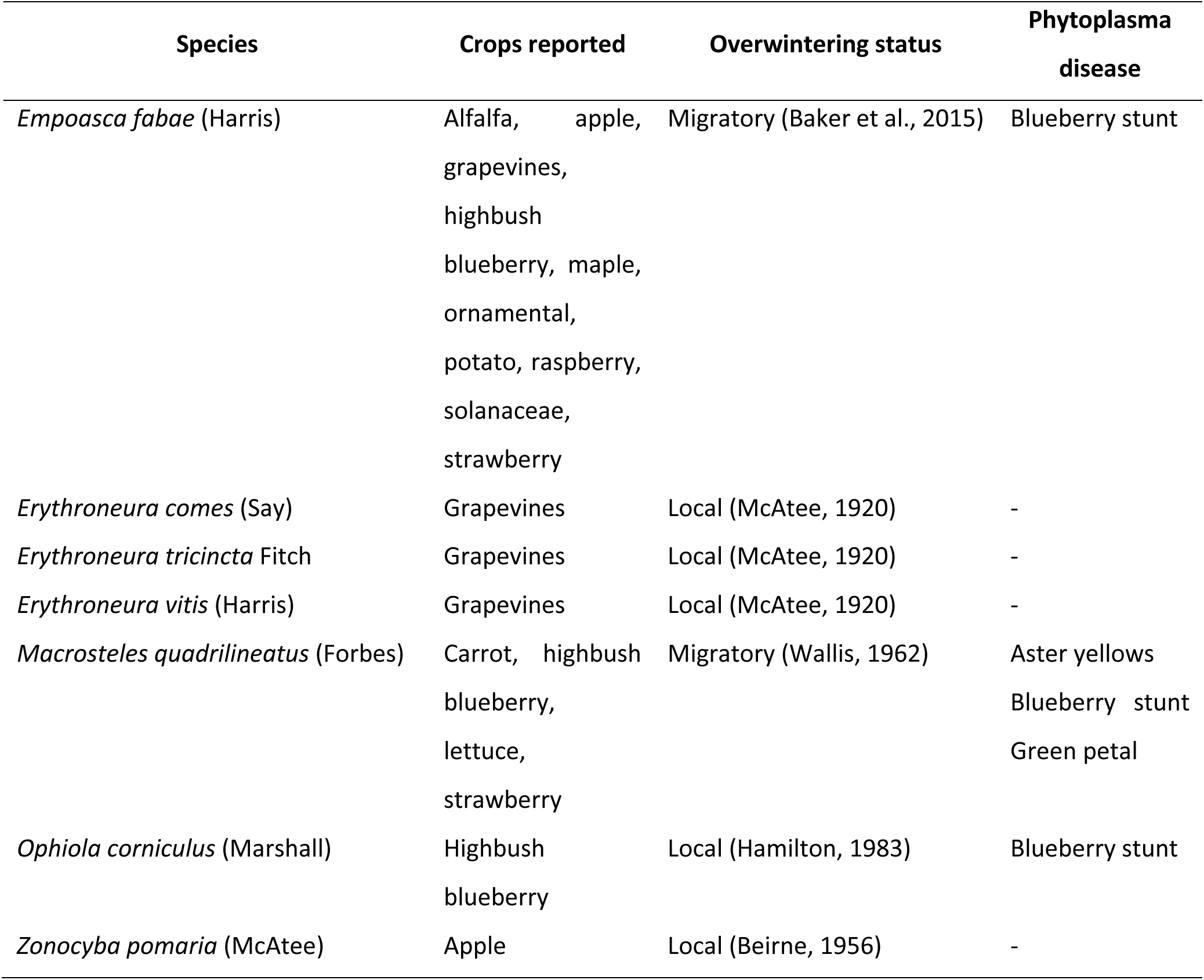
Leafhopper species and associated phytoplasma disease mentioned in the RAP network (2004-2025) in crops in Québec (Canada).

Within the RAP network, three phytoplasma diseases were linked to crops: aster yellows in carrot and lettuce, blueberry stunt in highbush blueberry, and green petal in strawberry. All three were associated with *M. quadrilineatus*, while *E. fabae* and *O. corniculus* were also linked to blueberry stunt (**Table 2**).

### Historical reports

Historical documents show that leafhoppers were first recorded in association with crops as early as 1886 for local species, and 1908 for migratory (*E. fabae*) (**Figure 4**, **Supporting Table S2**). Most species reported in historical documents also appeared in the RAP network, except for seven species: *Aceratagallia sanguinolenta* (Provancher), *Aphrodes bicincta* (Schrank), *Colladonus clitellarius* (Say), *Dikraneura mali* (Provancher), *Edwardsiana rosae* (Linnaeus), *Endria inimicus* (Say), and *Kyboasca maligna* (Walsh) (**Table 3**).

**FIGURE 4.**
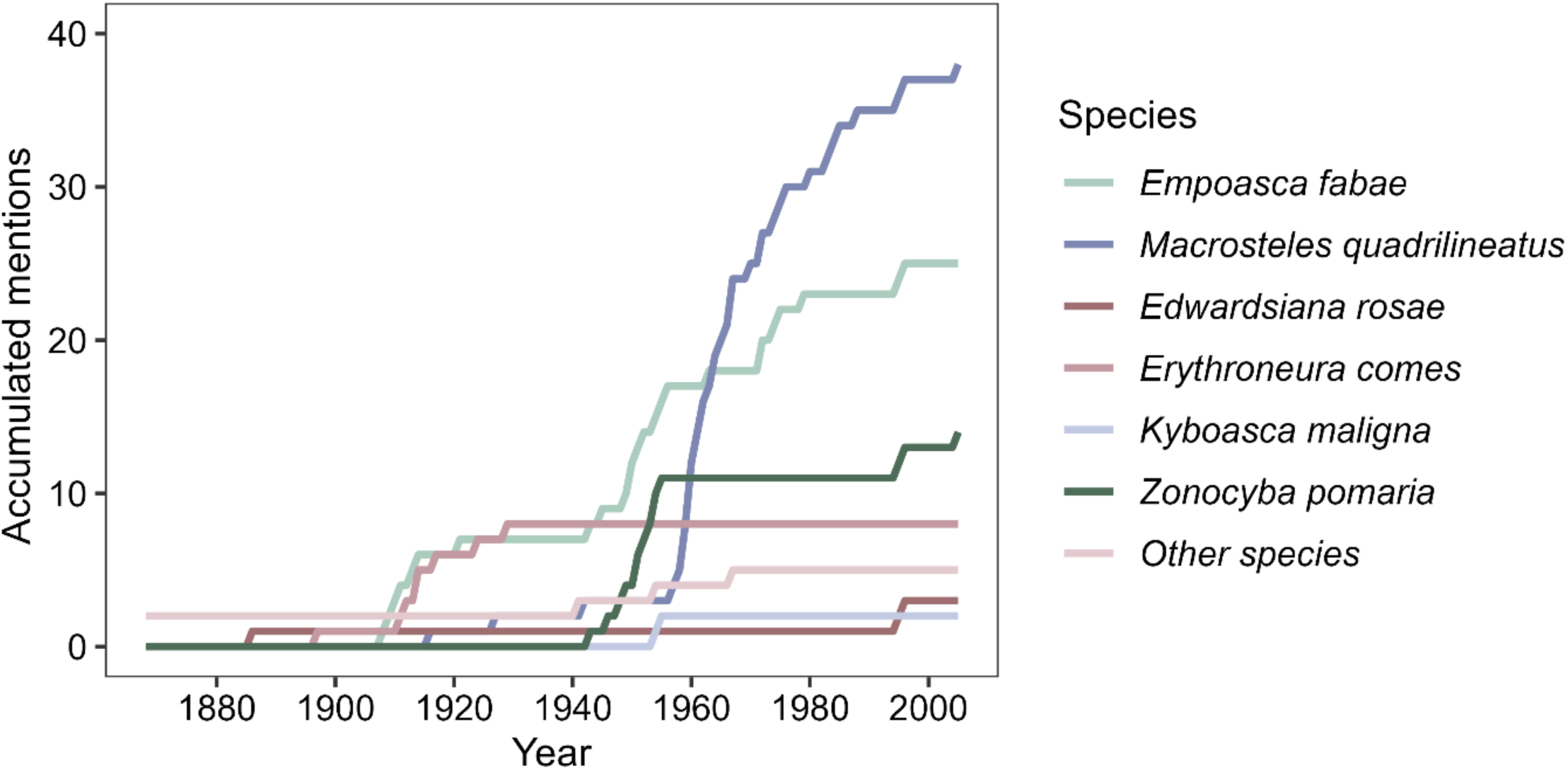
Accumulated historical records of field reports on leafhopper species linked to crops in Québec (Canada). References for these reports are listed in Supporting Table S2. Other species refer to those mentioned only once (*Aceratagallia sanguinolenta*, *Aphrodes bicincta*, *Colladonus clitellarius*, and *Dikraneura mali*).

**TABLE 3.** Leafhopper species and associated crops mentioned in historical documents in Québec (Canada). The accumulated mentions over time are displayed in Figure 4, and all references are provided in the Supporting Table S2.

| Species | Crops reported | Overwintering status | Disease |
| --- | --- | --- | --- |
| *Aceratagallia sanguinolenta* (Provancher) | Potato | Local (Hamilton, 1998) | Yellow dwarf |
| *Aphrodes bicincta* (Schrank) | Strawberry | Local (Chiykowski, 1970) | Green petal |
| *Colladonus clitellarius* (Say) | Peach | Local (Beirne, 1956) | - |
| *Dikraneura mali* (Provancher) | Apple | Local (Hamilton, 1983) | - |
| *Edwardsiana rosae* (Linnaeus) | Grapevines and roses | Local (Hamilton, 1983) | - |
| *Empoasca fabae* (Harris) | Alfalfa, apple, currant, grapevines, potato | Migratory (Baker et al., 2015) | - |
| *Endria inimicus* (Say) | Wheat | Local (Beirne, 1956) | - |
| *Erythroneura comes* (Say) | Grapevines | Local (McAtee, 1920) | - |
| *Kyboasca maligna* (Walsh) | Apple | Local (Beirne, 1956) | - |
| *Macrosteles quadrilineatus* (Forbes) | Beet, carrot, celery, Chinese asters, lettuce, potato, strawberry | Migratory (Wallis, 1962) | Aster yellow, green petals |
| *Zonocyba pomaria* (McAtee) | Apple, grapevines | Local (Beirne, 1956) | - |

Throughout the historical period, with mentions from 1886 to 2005, *M. quadrilineatus* received the most of them, followed by *E. fabae* and *Z. pomaria* (**Figure 4**). As in the RAP network, local species were primarily associated with a single crop (except *E. rosae* and *Z. pomaria*), whereas migratory species were associated with multiple crops (**Table 3**). Mentions of *M. quadrilineatus* were driven largely by its association with aster yellows phytoplasma in vegetable crops (carrot, celery, lettuce, onion) and green petal disease in strawberry. References for leafhopper–crop associations are available in **Supporting Table S2**.

### Museum records

Of the species documented in the RAP network and historical literature, ten had corresponding specimens in the Canadian National Collection (CNC), collected between 1906 and 2018. No specimens were found for *E. rosae*, *e. comes*, *O. corniculus*, or *Z. pomaria*. The earliest local species specimen was *E. vitis* (8 June 1906), followed by *D. mali* (24 May 1906) and *A. sanguinolenta* (5 May 1927). For migratory species, the earliest *E. fabae* specimen dates to 24 August 1914, with a spring specimen from 25 May 1950, while *M. quadrilineatus* first appeared on 20 June 1929, and a spring specimen on the 21^st^ of May 1951. Full specimen data, including all first and early collection dates, are provided in **Supporting Table S3**.

### Leafhopper assemblage on crops

We compiled available data on leafhopper assemblages in Québec crops spanning 1959–2024, including lettuce (1959–1964), alfalfa (1960), grapevines (1997–2008), highbush blueberry (2012), and strawberry (2021–2024) (Bostanian et al., 2003; Jacques et al., 2025; Lachance et al., 1961; Pagé, 2013; Perron & Crête, 1968; Plante et al., 2024; Saguez et al., 2014). The alfalfa and lettuce reports focused solely on *M. quadrilineatus*, representing a single-species population, but given the valuable historical context of its documented abundance, we included it.

In grapevines, local species were abundant, comprising up to 89% of specimens (**Figure 5**). In contrast, migratory species (*E. fabae* and *M. quadrilineatus*) accounted for over 50% of total abundance in highbush blueberry (2012) and strawberry (2021–2024), reaching a maximum of 88% in 2021 (**Figure 5**). In alfalfa and lettuce, where only *M. quadrilineatus* was surveyed, maximum counts reached 5,355 and 20,474 individuals, respectively, evidencing its historically high abundance in these crops.

**FIGURE 5.**
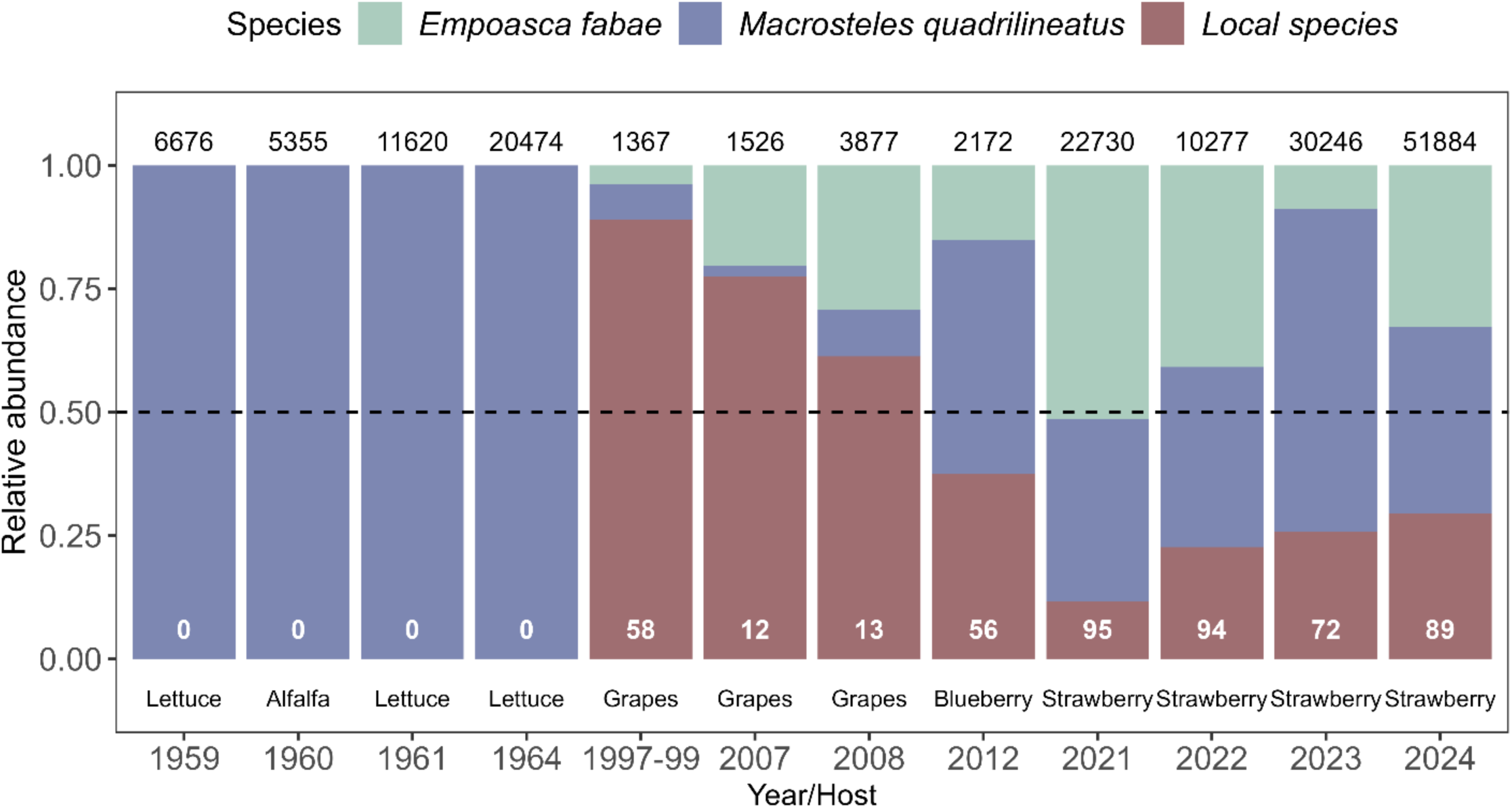
Proportional abundance of migratory versus local leafhopper species in published surveys from four crops in Québec (Canada). Each bar represents a different year/crop: alfalfa (1960), lettuce (1959, 1961, 1964), grapevines (1997–1999, 2007-2008), highbush blueberry (2012), and strawberry (2021–2024). Species are grouped into migratory (*Empoasca fabae* and *Macrosteles quadrilineatus*) and local groups. The number at the top of each bar indicates the total number of adults captured. Numbers within the local species bars indicate the number of species in this group. The dashed vertical line marks an equal proportion (0.5).

## DISCUSSION

Our findings indicate that the first field reports of leafhoppers in Québec are occurring earlier. However, this overall trend conceals considerable complexities. First reports advanced up to 3.5 days per year across carrot, lettuce, ornamental nurseries, potato, and strawberry. Nonetheless, these estimates should be interpreted with caution due to data limitations, including sparse records prior to the creation of the RAP network in 2004 and varying sample sizes across crops. Historical records reveal that leafhoppers have been recognized as pests since 1886, with mentions and perceived importance changing over time, and migratory species cited more frequently than local ones. Museum specimens confirm that species listed in the RAP network and in historical documents have been present in Québec for at least a century. Meanwhile, published surveys indicate an increase in the relative abundance of migratory species, although differences among crops are evident. Older surveys already documented high abundances of migratory species, suggesting their longstanding prominence and supporting a shift in leafhopper pest status in Québec, driven largely by crop-specific responses to migratory taxa. Given these caveats and observations, we now address the three questions outlined previously.

### Are first field reports of leafhoppers appearing earlier?

Before the RAP network was established in 2004, reports of leafhoppers were scarce, with only five from 1960 to 2000, making it difficult to assess trends during this period. After 2004, records became more frequent, but not immediately earlier, as a subtle decline appears to have started around 2013. If the increase in post-2004 RAP reports alone were driving earlier dates, we would expect a similar response across all crop categories. Instead, only five of the ten crops analyzed showed significant advances. These patterns suggest that while the RAP network improved pest reporting, it does not fully account for the observed advances in first-report dates.

### Do these early shifts indicate changes in pest status?

Multiple lines of evidence suggest that leafhopper pest status has shifted across crops in Québec, but the nature of this shift varies substantially by crop. Potato provides the clearest evidence. Widespread outbreaks of *E. fabae* in 2016 prompted expanded monitoring across seven regions of Québec in 2017 and the establishment of insecticide application thresholds (Martinez et al., 2022). Building on those intensive monitoring efforts, *E. fabae* pressure on potato has varied substantially over the past eight years, ranging from low (2018, 2022, and 2023) to moderate (2019) to high (2017, 2020, 2021, and 2024) (RAP, 2025). Our analysis shows that the first report of the potato leafhopper on potato fields advanced by two days per year from 2004 to 2025. This advance, occurring alongside documented pest pressure, suggests that factors beyond monitoring are driving earlier detection.

Strawberry presents a distinct shift, associated with phytoplasma disease and feeding injury in newly established fields. Strawberry green petal disease, a phytoplasma-related disease transmitted by leafhoppers, was frequently reported in the province between 1950 and 1970 (Santos et al., 2025b). Although this disease prompted extensive monitoring in Québec and the Atlantic provinces between 1971 and 1972, economic impacts were considered minimal at that time (Chiykowski et al., 1973). However, reports of phytoplasma disease have re-emerged across multiple crops in Québec over the past decade (Brochu et al., 2023), including green petal disease in strawberry (Plante et al., 2024). This has led to ongoing research to elucidate the role of leafhoppers as strawberry pests, their assemblage composition (Jacques et al., 2025; Plante et al., 2024), and the extent of the damage they may cause, is consistent with the rapid advance in first reports we observed for this crop.

Carrot, lettuce, and ornamental nurseries present less-developed evidence of pest status shifts. In carrot and lettuce, the migratory leafhoppers documented by the RAP (primarily *M. quadrilineatus*) are recognized as vectors of aster yellows phytoplasma (Olivier et al., 2009), and field reports of this pathosystem have been documented in the province since 1942 (Miller & De Lyzer, 1960). While this disease remains present in both crops in Québec (Brochu et al., 2023), the associated economic losses have yet to be determined. Ornamental nurseries present a more complex case, encompassing diverse plant species that may host multiple leafhopper taxa, including *E. fabae*, with varying phenologies and vector status, potentially reflecting multiple underlying causes that warrant further investigation.

The remaining five crop categories studied here, including alfalfa, apple, grapevines, raspberry, and solanaceous crops, showed no significant variation in first-report dates. In alfalfa, the change in pest status seems more recent, with outbreaks of *E. fabae* recorded between 2019 and 2021 (RAP, 2019, 2021), which may explain the marginal significance in our analysis (*p* = 0.088). Despite this short-term record, recent research indicates increasing concern about the status of this pest species in this crop. Shi et al. (2025) evaluated multiple control strategies for *E. fabae* in alfalfa, the first such study in Québec, to the authors’ knowledge. This emerging focus is notable because *E. fabae* has long been a major pest of alfalfa in the neighboring province of Ontario (Appleton et al., 2003). Such recent attention suggests that Québec may be experiencing similar pressures.

For apple, grapevines, raspberry, and solanaceous crops, three points may explain the absence of a significant trend. First, the leafhopper assemblage in most of these crops appears to be dominated by local species, with more consistent phenology. In grapevines, for instance, leafhopper assemblages consist mainly of local species (Bostanian et al., 2003, 2006; Saguez et al., 2014), although migratory species can be present (Vincent et al., 2018). Historically, managing leafhoppers has been less of a priority in these crops, with leafhoppers classified as a secondary pest with some importance (Vincent et al., 2018). The marginal trend observed in grapevines (p = 0.075) may reflect the detection of the invasive leafhopper *Euscelidius variegatus* (Kirschbaum) in Eastern Canada, which is known as a vector of multiple phytoplasmas in Europe, and could signal future shifts in pest status (Parent et al., 2019). Second, other invasive pests may have shifted monitoring priorities, as seen with the initial report of spotted wing drosophila in Québec in 2012 (Saguez et al., 2013). Third, although leafhoppers are vectors of multiple phytoplasma diseases in Québec (Almeida Santos et al., 2024; Olivier et al., 2009), the prevalence of these diseases appears limited in many crops (Brochu et al., 2023). Even after the first detection of globally important diseases in Canada, i.e., *Bois noir* (Rott et al., 2007), the maximum rate of phytoplasma-infected leafhoppers in grapevines was 8.3% (Olivier et al., 2014), suggesting that phytoplasma diseases remained below economically relevant levels, but call for continuous surveillance (Carisse et al., 2017).

Importantly, the vector status of leafhoppers amplifies the surveillance effect. When a leafhopper species is known to transmit diseases or when infected plants are observed in the field, the incentive to monitor intensifies (Duffeck et al., 2025). An infected leafhopper can threaten crop production, making early detection not just informative but, to some extent, economically protective (Ben Moussa et al., 2020; Parent et al., 2019). The presence of a known disease vector or disease symptoms justifies monitoring efforts that might otherwise seem disproportionate to the insect’s direct feeding damage (Carisse et al., 2017).

However, increased surveillance of phytoplasma diseases in Québec has improved detection without necessarily indicating economic impact (Brochu et al., 2023). Reports of aster yellows, green petal disease, and blueberry stunt have become more frequent, but this likely reflects greater perception and monitoring effort rather than increased crop losses. This distinction is particularly important when considering the crop cycle. In short-cycle crops, symptoms of phytoplasma diseases may appear late in the season, affecting harvestable yield and making economic impacts less apparent (Olivier et al., 2009). In contrast, perennial crops (e.g., grapevines and highbush blueberries) can accumulate infections over years, potentially leading to more severe long-term consequences even if annual detection rates remain low (Arocha-Rosete et al., 2019; Olivier et al., 2014). Thus, increased phytoplasma detections do not certainly translate to economic damage, and threshold-based management remains essential.

### Are these changes on pest status associated with migratory species, local species, or both?

Shifts in leafhopper species composition across crops offer additional evidence for the patterns observed in first-report dates. The crops showing significant advances (potato, strawberry, carrot, lettuce, and ornamental nurseries) are those in which migratory species are reported to be abundant. In contrast, the crops showing no significant trend (apple, grapevines, raspberry, solanaceous crops) are dominated by local species, with more consistent phenology and historically less economic concern. This pattern strongly suggests that the observed advances are driven by migratory species.

This interpretation is supported by documented phenological advances in both key migratory species. For *E. fabae*, first report dates in North America have advanced by approximately 10 days over the past six decades (Baker et al., 2015), and in Québec, first adult captures occur earlier than degree-day models predict, suggesting that warming, wind patterns, and host availability interact to advance its phenology (Santos et al., 2025). Suction trap data from the US Midwest further show that *E. fabae* activity is extending later into the year, potentially broadening its window of economic impact (Lagos-Kutz et al., 2024). For *M. quadrilineatus*, while quantitative phenology data are sparser, its well-documented migratory behavior and role as the primary vector of aster yellows (Olivier et al., 2009; Romero et al., 2026) suggest it may respond similarly to climatic drivers (Plante et al., 2024).

These documented advances have a direct mechanistic basis. Migratory species, whose arrival timing depends on continental wind patterns and source population phenology further south, are likely more responsive to climate variability than local species whose phenology is governed by local overwintering conditions (Plante et al., 2024). The increasing proportion of migrant leafhoppers in crop assemblages could thus intensify any climate-driven signals in first-report dates, particularly for crops where these species are most abundant, such as potato and strawberry. Together, these lines of evidence converge on migratory species as the principal drivers of the shifts we observed, while also pointing to the need for continuous monitoring as climate paradigms continue to change.

## CONCLUDING REMARKS

Our results indicate that first field reports of leafhoppers are occurring earlier across five crops, while also supporting the historical and continued prominence of migratory species in Québec crop pest assemblages. These advances are unlikely to reflect monitoring effort alone, since migratory species have been documented in Québec for over a century, and their relative abundance has increased in recent surveys. Rather, improved surveillance, increasing migratory pest pressure, the apparent resurgence of phytoplasma diseases, and climate-driven phenological shifts have elevated the economic importance of leafhoppers in Québec agriculture. Thus, the main change may be as much in our perception and management of these insects as in the insects themselves.

Whether this renewed attention will persist, or whether leafhoppers will once again recede from concern during quieter years, remains an open question. The historical record suggests that such oscillations are possible, as periods of heightened concern have alternated with periods of relative neglect throughout the past century. However, under projected warming scenarios, leafhopper pest species are expected to remain a consistent and potentially intensifying presence in Eastern Canada (Santos et al., 2024), making continuous monitoring not just useful but necessary. What was once ‘*loin des yeux, loin du cœur*’ (far from the eyes, far from the heart) appears to be consistently settling into ‘*près des yeux, près du cœur*’ (close to the eyes, close to the heart), and the evidence suggests it may stay that way.

## AUTHOR CONTRIBUTIONS

**Abraão Almeida Santos**: Conceptualization; data curation; formal analysis; investigation; methodology; visualization; writing – original draft; writing – review and editing. **Edel Pérez-López**: Resources; writing – review & editing.

## ACKNOWLEDGMENTS

We thank the *Réseau d’avertissements phytosanitaires* (RAP-MAPAQ) for developing and maintaining a comprehensive, publicly accessible phytosanitary surveillance network in Québec. We also acknowledge the contributions of the RAP network contributors (agronomists, crop advisors, farmers, and journalists), whose collaborative efforts enable these surveillance reports and offer a valuable resource for understanding pest and plant pathogen dynamics in Québec. Access to digitized specimens from the Canadian National Collection of Insects, Arachnids, and Nematodes (CNC) and the multiple libraries that preserve important historical resources were essential for establishing the historical context of this study. We also thank Valérie Boulva for allowing access to the Provancher Hemiptera collection at Université Laval. The graphical abstract was created in part using images from Servier Medical Art (https://smart.servier.com, CC BY 4.0 license) and BioRender (https://www.biorender.com/). We acknowledge the use of Grammarly (v. 1.2.250.1876) to assist with the language fluency and readability of this manuscript. All content was reviewed and approved by the authors, who take full responsibility for the accuracy and integrity of the work.

## FUNDING INFORMATION

This work was funded by the Réseau québécois de recherche en agriculture durable (RQRAD), the Ministère de l’Agriculture, des Pêcheries et de l’Alimentation du Québec (MAPAQ), and the Fonds de recherche du Québec–Nature et technologies (FRQNT) through the Programme de recherche en partenariat—Agriculture durable—Volet II—2e concours (application number 337847), as well as by the Natural Sciences and Engineering Research Council of Canada (NSERC) through the Alliance-SARI Program (Grant ALLRP 588519-23). EPL also thanks the CRC program for its support.

## CONFLICT OF INTEREST STATEMENT

The authors are not aware of any conflicts of interest associated with this research.

## DATA AVAILABILITY STATEMENT

All data supporting the findings of this study are available in the supporting information.

## SUPPORTING INFORMATION

**FIGURE S1.** Diagnostic plots for the generalized additive mixed model analyzing trends in first leafhopper reports (1983–2025). Plots show normalized residuals after adjusting for the first-order autoregressive [AR(1)] correlation structure. From top left to bottom right: Normal Q-Q plot with a reference line; Residuals versus fitted values with a horizontal line at zero; Histogram with an overlaid normal density curve; and Autocorrelation function (ACF) of normalized residuals.

**Material S1.** Historical documents search and associated online sources.

**Table S1.** First field observation per crop per year in 16 crops in Québec (1960-2025). The Julian day (day of the year) and the report source are indicated.

**Table S2.** Historical mentions (1886-2005) of leafhoppers and associated crops, with references for reports in Québec.

**Table S3.** Leafhopper specimen records for species mentioned in the RAP network and historical documents obtained from the Canadian National Collection of Insects, Arachnids, and Nematodes (CNC, Ottawa) through the collection’s online portal (https://cnc.agr.gc.ca/taxonomy/TaxonMain.php).

## Notes

### Competing Interest Statement

The authors have declared no competing interest.

