## Supplementary material for "Leafhoppers’ pest status in Québec from 1868 to 2025": Figure S1

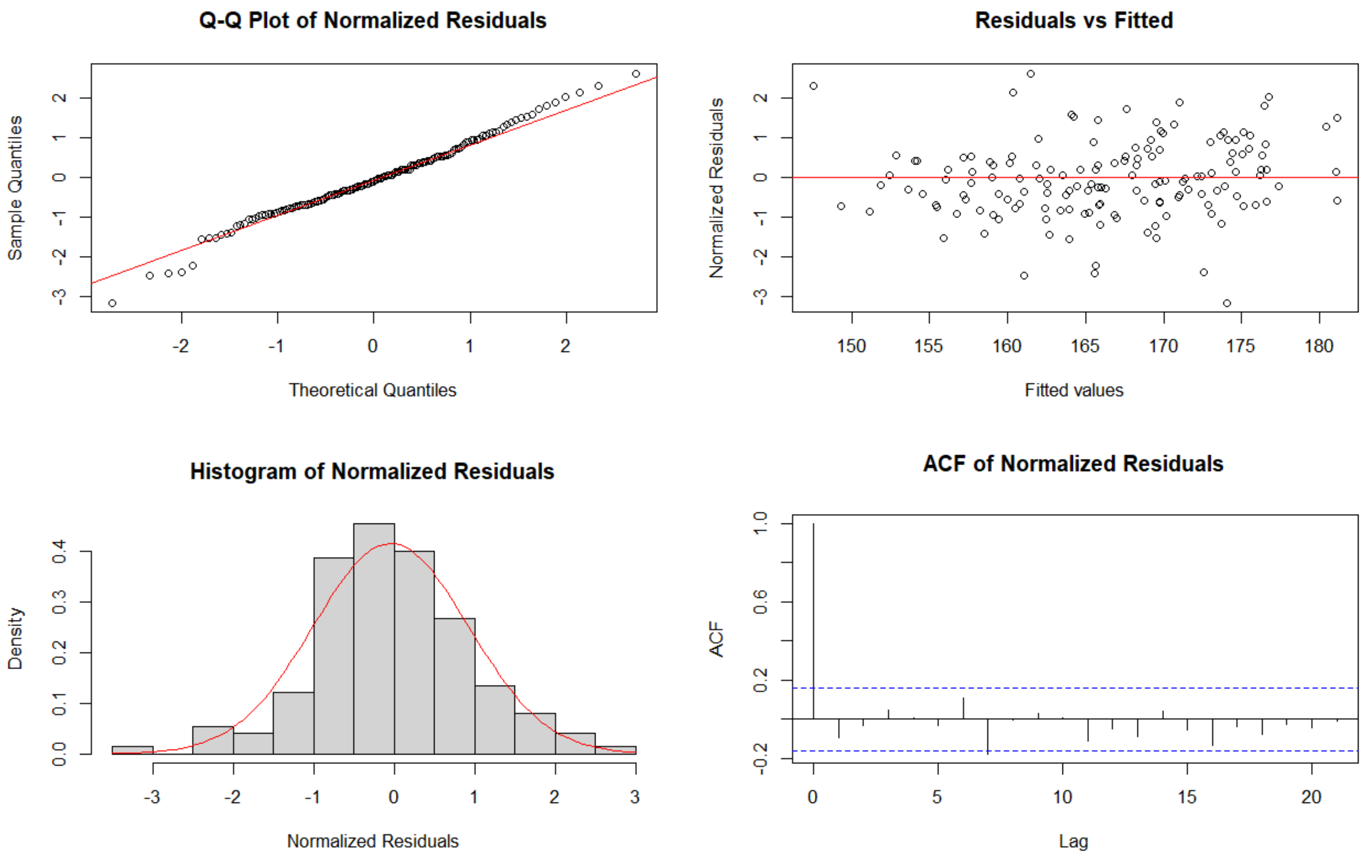


**SUPPORTING FIGURE S1** Diagnostic plots for the Generalized Additive Mixed Model examining trends in first leafhopper reports (1983–2025). Plots are based on normalized residuals after accounting for the first-order autoregressive [AR(1)] correlation structure. Normal Q-Q plot with reference line. Residuals versus fitted values with horizontal line at zero. Histogram with overlaid normal density curve. Autocorrelation function (ACF) of normalized residuals. As expected, the ACF shows a spike at lag 0 (exact autocorrelation), with no significant lags thereafter. Model diagnostics indicated no significant deviation from normality (Shapiro-Wilk test: $p = 0.588$) and no significant residual autocorrelation (Box-Ljung test: lag 5, $\chi^2 = 6.04$, $p = 0.302$; lag 10, $\chi^2 = 14.19$, $p = 0.165$).
