## Supplementary Material for "Leafhoppers’ pest status in Québec from 1868 to 2025"

**SUPPORTING MATERIAL S1**

**Historical documentation**

Because historical documents are not indexed in electronic databases and many exist only in print, we conducted targeted archival searches using the Biodiversity Heritage Library (BHL; https://www.biodiversitylibrary.org/) and other sources to trace associations between leafhopper pest species and crop field reports in Québec. Although these documents lack the precise timing needed to analyze the first reports, they provide context on the economic importance of leafhoppers over time.

For each document, we initially reviewed the title, abstract, and species index when available. We then searched for French and English keywords related to leafhoppers, phytoplasma diseases, and associated terms: aster yellow, cicadelle(s), leaf-hopper(s), leafhopper(s), hopper(s), green petal, jaunisse de l'aster, phytoplasmas, virus-like, yellows, tip-burn, and hopperburn. When mentions were found, we extracted the year, pest species, associated crops, and diseases. All leafhopper species names were verified for synonyms using the World Auchenorrhyncha database.

*Le Naturaliste Canadien*, one of the oldest journals documenting the natural history of Québec, has been published continuously since 1868 and was examined across all issues from 1868 to 2025 via the journal website (https://www.provancher.org/publications/le-naturaliste-canadien-3/). We also consulted the Provancher Hemiptera checklist, available through BHL (Provancher, 1886), which provides the initial formal description of leafhopper species in Québec, based on publications in Le Naturaliste Canadien. Species names in this checklist were later revised to account for synonyms (Duzee, 1912).

The Journal of the Entomological Society of Ontario, originally titled the Annual Report of the Entomological Society of Ontario and later the Proceedings of the Entomological Society of Ontario, was first published in 1871. It is the earliest journal to document pest reports from all Canadian provinces, including annual summaries of crop-affected species from the previous year. We examined all issues using the BHL database (1871–2001) and the journal website (2002–2025; https://www.entsocont.ca/archives.html).

The *Société d'entomologie du Québec* has published continuously since 1956 under three successive titles: the *Annales de la Société entomologique du Québec* (1955/56–1983), the *Revue d'entomologie du Québec* (1984–1992), and the bulletin Antennae (1994–present). The first two publications are available only in print and were consulted by the first author at the Université Laval library in February and March 2026. Antennae issues were accessed through the Bibliothèque et Archives nationales du Québec website (https://numerique.banq.qc.ca/).

We also reviewed the *Rapport annuel de la Société de Québec pour la protection des plantes* (1908–1961) via the Bibliothèque Cécile-Rouleau (https://bibliotheque.cecile-rouleau.gouv.qc.ca/documents/archives/pgq/A38A1_S633/), as well as its current version, the journal Phytoprotection, through printed volumes at the Université Laval library (1963–1989) and online volumes (1990–2025) available on the Érudit website (https://www.erudit.org/fr/revues/phyto/#back-issues). Additionally, we examined the reports from the *Rapport annuel de la Société de Pomologie et de Culture Fruitière* (1894–1960) via the Bibliothèque Cécile-Rouleau (https://bibliotheque.cecile-rouleau.gouv.qc.ca/).

Other sources consulted include A Synopsis of Economic Entomology (Lochhead, 1914) and *Les principales espèces d'insectes nuisibles et les maladies végétales* (Huard, 1916), both accessed through BHL. Finally, we reviewed the bulletin Nouv'Ailes (1999–2023), available on the website of *l'Association des entomologistes amateurs du Québec* (https://www.aeaq.ca/bulletins-nouvailes), and the Canadian Plant Disease Survey Archive (1920–2025; https://phytopath.ca/publications/canadian-plant-disease-survey-archive/).
